# Oncogenic KRAS regulates secretion of extracellular vesicles and surface membrane charge via regulation of phosphatidylserine in pancreatic cancer

**DOI:** 10.1101/2025.09.21.677641

**Authors:** Kshipra S Kapoor, Xin Luo, Kaira A Church, Martin M Bell, Bo Fan, Seoyun Kong, Elena V Ramirez, Yi-Lin Chen, Fernanda G Kugeratski, Sibani Lisa Biswal, Kathleen M McAndrews, Florian Gebauer, Christoph Kahlert, Jacob T Robinson, Raghu Kalluri

## Abstract

Extracellular vesicles (EVs) have the potential to be used as a liquid biopsy for cancer detection and treatment response assessment. Although the potential of EVs as disease-specific biomarkers has been promising, the rapid and specific enrichment of EVs from body fluids in the clinical setting remains challenging. To address this limitation, we developed a Microfluidic Electrophoresis (MEP) device, that allows label-free enrichment of EVs based on charge from the serum of patients with pancreatic cancer (PaCa). The ability of MEP to enrich anionic EVs was validated using cell line-derived EVs and PaCa serum-derived EVs. We observed a positive correlation between the KRAS status, secretion of EVs, and net negative zeta potential (ζ-potential) of EVs. Further analyses identified phosphatidylserine (PS) and extraluminal DNA as molecular determinants of the enhanced anionic nature of PaCa-derived EVs. Overall, this work introduces a new microfluidic device to enrich cancer EVs in circulation with potential for further rapid detection of PaCa.

## Introduction

Pancreatic Cancer (PaCa) is currently the fourth leading cause of death due to malignancy and is expected to be ranked second by 2030^1, 2^. About 90% of PaCa is pancreatic ductal adenocarcinoma (PDAC), for which KRAS mutations are the primary drivers^3^. KRAS mutations are frequently found in codon 12, with the most common being G12D, trailed by G12V^4, 5^. Lack of early detection, metastatic disease, and limited treatment options contribute to the high mortality rate of PaCa^4–6^. Therefore, it is imperative that we attempt to develop rapid and sensitive strategies for screening PaCa patients at early stages with the potential for enhanced benefit from available treatments, thereby reducing PaCa-associated mortality.

An emerging approach for early diagnosis of PaCa is based on the use of liquid biopsies, which are simpler to procure than tissue biopsies employing minimally invasive methods. The possibility of employing EVs in the serum as cancer liquid biopsy has recently attracted attention^7–13^. Exosomes (40-150nm diameter), a subtype of EVs, are endosomal-derived vesicles with a phospholipid bilayer membrane that possess diverse molecular cargoes and mediate intercellular communication^12, 14–17^. Additionally, EVs may play a role in cancer progression and metastasis, cellular homeostasis, innate and adaptive immunity, and angiogenesis^18–22^. Their unique biology such as their lipid bilayer envelope and the presence of an array of biomolecules on the surface of their lumen^23^, make them a promising candidate as a potential disease diagnostic. Circulating EVs comprise both tumorigenic EVs and non-tumorigenic EVs (non-EVs). Unfortunately, the latter can dilute with the genomic assessment of EVs and can compromise sensitivity and specificity^24^. Therefore, methods to enrich tumorigenic EVs to meet the detection threshold required by most downstream analysis are essential to facilitate early recognition of PaCa^25^. However, selectively enriching cancer-derived EVs from serum or plasma remains challenging^26,27^.

Routinely used EV isolation techniques include ultracentrifugation, size exclusion, density gradient, and precipitation methods, isolate all EVs in a sample^28^. Alternative approaches that use silica substrates and antibody-coated beads to isolate EVs have been minimally explored^29–31^. The antibody-coated beads assays involve multiple labelling steps, are time consuming and still cannot differentiate between cancer EV versus non-cancer EVs^32, 33^. Additionally, identifying critical molecular biomarkers on the EV surface that are specific to cancer cells remains a challenge^31, 34–36^. Such efforts have also been further hindered by EV heterogeneity within individual patient samples^37, 38^, complicating specificity determination.

To address this challenge, we developed a new label-free microfluidic device - Microfluidic Electrophoresis (MEP) platform that utilizes an electric field to enrich EVs from the serum of PaCa patients based on the EV surface charge (zeta potential [ζ-potential]), which we discovered to be a hallmark of PaCa EVs. Additionally, we reported a link between oncogenic Kras expression, EV production and negative surface ζ-potential. Furthermore, we discovered that extraluminal DNA and the anionic phospholipid Phosphatidylserine (PS) on the EV surface contributes to the net negative ζ-potential of PaCa EVs, which is influenced by oncogenic Kras. Overall, these results demonstrate that MEP platform can be used as a tool for the enrichment of cancer EVs from the serum of patients with cancer, with potential for early detection of PaCa, by exploiting the anionic charge of PaCa EVs at its basic biophysical property.

## Results

### Oncogenic KRAS regulates EV production and net charge

We evaluated the molecular and biophysical characteristics of EVs isolated from seven different cell lines of pancreatic origin: human non-malignant epithelial cell lines (HPNE, HPDE, HK-2) and human malignant PDAC cell lines (Panc-1, T3M4, MiaPaCa-2, and BxPC3). Nanoparticle Tracking Analysis (NTA) showed that the isolated nanoparticles had a size distribution typical of small EVs (**Supplementary Figure 1A**). Cryogenic electron microscopy (Cryo-EM) analysis on both tumorigenic (Panc-1) and non-tumorigenic (HPNE) EVs demonstrated that the isolated particles were enclosed by a lipid bilayer (**Supplementary Figure 1B**). To further extend the characterization of EVs, we employed the bead-based flow cytometry approach to measure the expression levels of the tetraspanins, CD9, CD63, and CD81, which are commonly used as EV biomarkers, and demonstrated the presence of these markers from all the cell lines examined (**Supplementary Figure 1C**).

The surface charge of EVs was evaluated by determining the ζ-potential of EVs derived from the pancreatic cells. ζ-potential is an indicator of a membrane’s net charge and is representative of the particle charge in relation to its surrounding medium^39^. In the non-tumorigenic and tumorigenic pancreatic lines, we observed a positive correlation between negative EV ζ-potential and increased aggressiveness of pancreatic cells (**Figure 1A-B**). The aggressiveness of the pancreatic cell lines was determined using a soft agar colony formation assay (**Supplementary Figure 1D**) and correlated with the EV charge (**Figure 1B**). Since the mutant form of KRAS is the key driver of pancreatic cancer, we used ELISA to measure the RAS activity in tumorigenic pancreatic cell lines (Panc-1 and BxPC3) and non-tumorigenic pancreatic epithelial cell line (HPNE) and observed a positive correlation of R^2^=0.8788 between activated RAS levels and the negative ζ-potential of EVs derived from these cell lines (**Figure 1C-D**).

**Figure 1:**
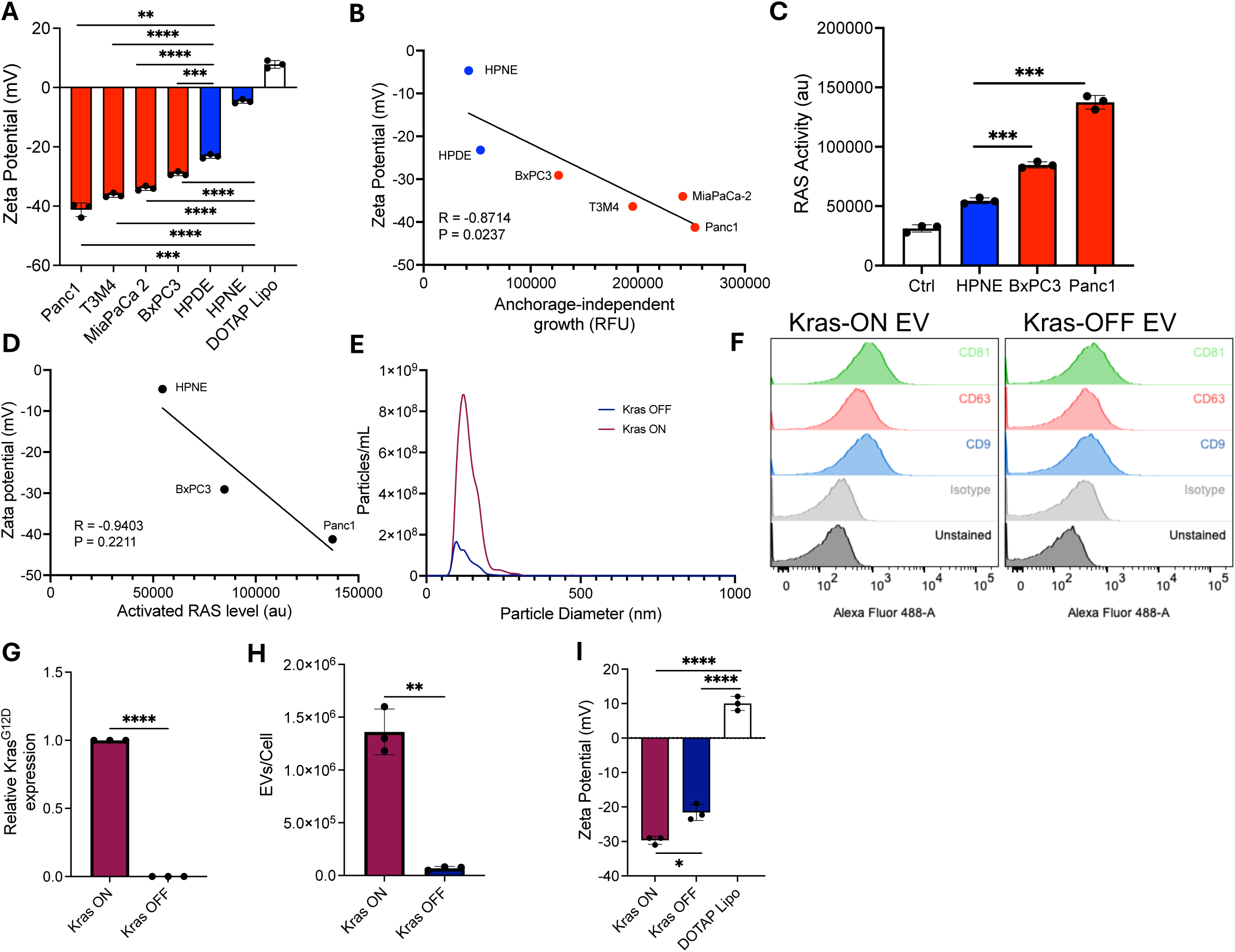
Oncogenic Kras regulates EV production and surface charge. **(A)** ζ-potential of EVs released by the indicated tumorigenic and non-tumorigenic pancreatic cell lines. **(B)** The correlation between negative value of ζ-potential and aggressiveness tested in soft agar assay evaluated using Pearson correlation. **(C)** Kras status in the respective cell lines measured by RT-qPCR analysis. **(D)** Measurement of RAS activity and correlation between activated RAS levels and ζ-potential of EVs from respective cell lines. **(F)** The detection of surface marker CD9, CD63, and CD81 on EVs by flow cytometry before and after the induction of Kras knock down in parental cells. **(G)** The induction of Kras^G12D^ in EVs from U785 cells, confirmed by RT-qPCR analysis. **(H)** EV production from parental cells before and after Kras knock down. **(I)** ζ-potential of EVs before and after Kras knock down in parental cells. DOTAP liposomes used as a control. Data is represented as Mean ± SD. Statistical significance was determined using Unpaired t-test or Pearson Coefficient (∗p ≤ 0.05, ∗∗p ≤ 0.01, ∗∗∗p ≤ 0.001, ∗∗∗∗p ≤ 0.0001,) n = 3 biological replicates.

To gather additional evidence that Ras activity might directly influence the overall net negative charge of EVs, we utilized an inducible oncogenic Kras expression system to evaluate the role of mutant Kras^G12D^ (doxycycline inducible Kras^G12D^ On, and Kras^G12D^ Off [iKras cell line]) in EV secretion and contribution to EV surface charge (**Figure 1E-I**). NTA analysis revealed that isolated EVs from both Kras-on and Kras-off cells had a size distribution typical of small EVs and expressed canonical-EV tetraspanins (**Figure 1E-F, Supplementary Figure 1A**). Induction of KRAS^G12D^ was validated by RT-qPCR (**Figure 1G**) and KRAS^G12D^ induction was associated with a significant increase in EV secretion (**Figure 1H**). Additionally, EVs derived from Kras-on cells were significantly more anionic than EVs derived from Kras-off cells (**Figure 1I**).

### Contribution of distinct biomolecules towards net charge of EVs

To determine the role of distinct biomolecules that could contribute to the charge of EVs, we devised approaches to evaluate the contribution of extraluminal DNA, surface proteins, and lipids in determining the anionic charge of EVs.

To evaluate whether surface proteins are linked to the negative charge of EVs, we cleaved the extracellular domain of membrane proteins on the surface of EVs by using Proteinase-K (Pro-K) (**Figure 2A-D**). NTA analysis shows that Pro-K treatment of EVs did not induce major alterations in their size distribution or their concentration (**Figure 2A**). Additionally, we tested whether Pro-K treatment was successful at digesting the majority of extracellular domains of membrane proteins using flow cytometry of bead-bound EVs. Using the tetraspanin CD81 as a readout, we observed that expression of CD81 was decreased upon Pro-K treatment compared to untreated EVs (**Figure 2B**). We also observed that Pro-K treated and untreated EVs from both tumorigenic Panc1 cells and non-tumorigenic HPNE cells had equivalent negative charges with no significant difference in ζ-potential (**Figure 2C**). To determine the integrity of EVs after Pro-K treatment, we quantified the levels of protein localized in the lumen of EVs. We used single particle flow analysis to detect GFP localized in the intraluminal space by labeling Panc1 EVs with CD63-GFP labelled Panc1 EVs (**Figure 2D**). GFP levels of Pro-K treated and untreated samples revealed no differences in GFP levels between the samples, supporting the fact that the integrity of lipid bilayer was not disrupted (**Figure 2D**). These findings suggest the proteins on the EV surface, do not alter the EV surface charge.

**Figure 2:**
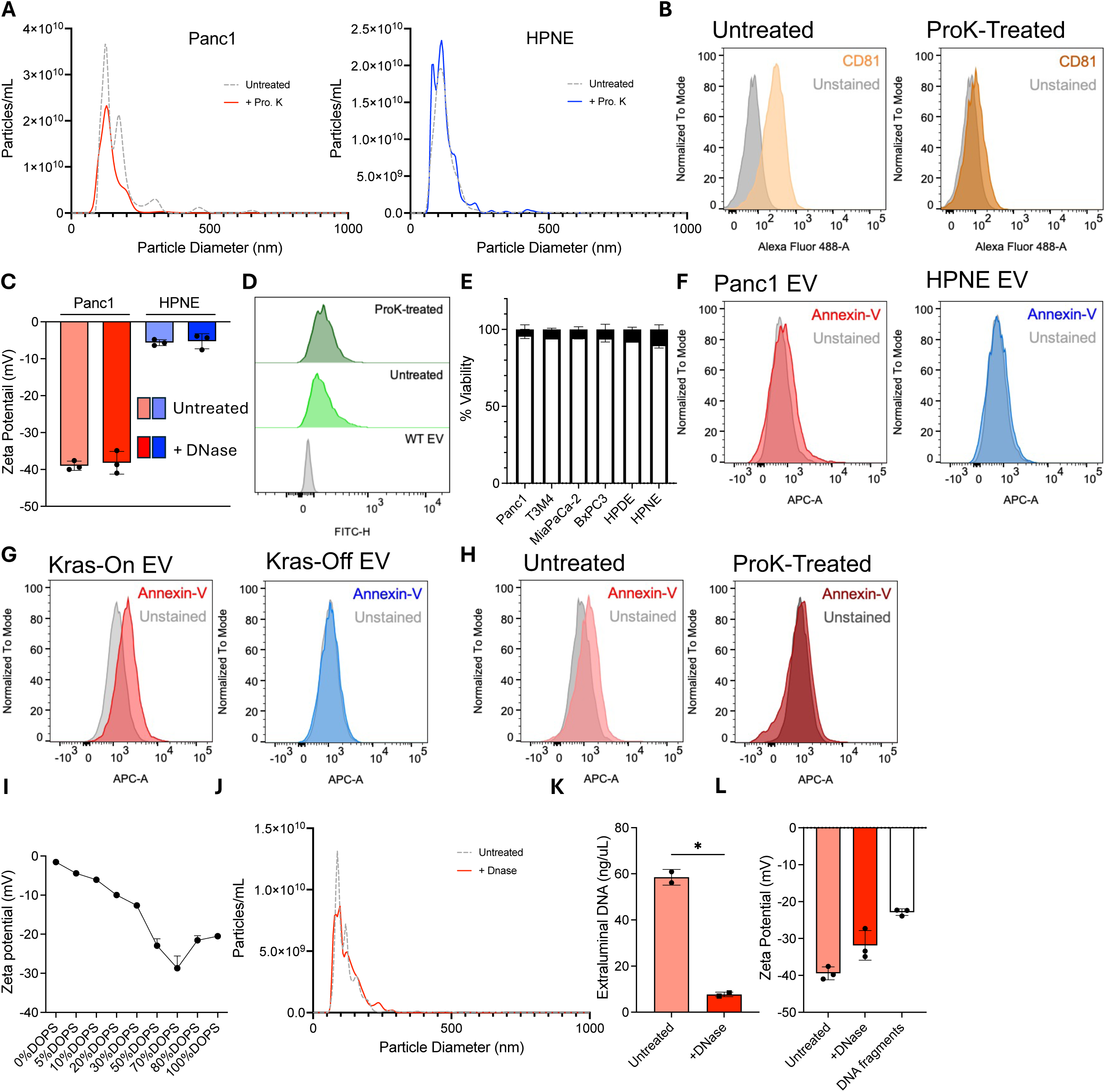
Contribution of distinct biomolecules to EVs charge. **(A)** Size characterization before and after the Proteinase K treatment for EVs derived from tumorigenic (Panc1) and non-tumorigenic (HPNE) cells. **(B)** The efficiency of proteolysis was determined by measurement of surface CD81 of EVs derived from Panc-1 cells by flow cytometry. **(C)** ζ-potential of the untreated and proteinase K treated EVs quantified by Malvern Zetasizer. **(D)** Representative histograms of the detection of EV luminal GFP proteins fused with CD63 with or without Proteinase K treatment by flow cytometry. **(E)** Viability of serum starved cells accessed by trypan blue assay. **(F)** Detection of surface PS level on EVs from tumorigenic (Panc1) and non-tumorigenic (HPNE) cells, by flow cytometry, indicated by the staining of PS with Annexin-V. **(G)** Detection of surface PS level on untreated and proteinase K treated EVs from Panc1 cells, by flow cytometry, indicated by the staining of PS with Annexin-V. **(H)** Detection of surface PS level on EVs from U785 cells before and after the induction of Kras^G12D^, by flow cytometry, indicated by the staining of PS with Annexin-V. **(I)** Detection of ζ-potential on synthetically fabricated liposomes with variable levels of PS. **(J)** Size characterization before and after the DNase treatment for EVs derived from Panc1 cells. **(K)** Quantification of cell free/extraluminal DNA before and after DNase treatment. **(L)** ζ-potential of the untreated and DNase treated EVs quantified by Malvern Zetasizer. Data are represented as Mean ± SD. Statistical significance was determined using Unpaired t-test (∗p ≤ 0.05, ∗∗p ≤ 0.01, ∗∗∗p ≤ 0.001, ∗∗∗∗p ≤ 0.0001,) n = 3 biological replicates.

To evaluate whether anionic phospholipid Phosphatidylserine (PS) contributes to the negative EV charge, we used flow cytometry to evaluate the PS levels on the EV surface by measuring the PS-binding protein, annexin V (**Figure 2E-I**). PS levels are associated with cell death, therefore, as a control we employed a trypan blue assay to determine the presence of dead cells in the cells used for EV isolation. We found low levels of cell death in the cell lines used for EV production in this setting (**Figure 2E**). The PS positivity of tumorigenic Panc1 cell derived EVs was significantly higher compared to that of non-tumorigenic cell derived EVs (**Figure 2F**). A high degree of positive correlation was observed between the PS levels and anionic ζ-potential of EVs (**Figure 2F**). Taken together, these results indicate that, the PS levels on the surface of EVs secreted from tumorigenic pancreatic cells were significantly higher compared to EVs secreted from non-tumorigenic cells.

Earlier, we demonstrated that EVs derived from mutant iKras cell line (oncogenic Kras-on) possessed an overall negative ζ-potential compared to EVs derived from oncogenic Kras-off cells. Therefore, we next evaluated whether the PS level on these EVs varied depending on their oncogenic Kras status. We observed that the EVs extracted from oncogenic Kras-on cells had significantly higher PS level than those from oncogenic Kras-off cells indicating that activated oncogenic Kras helps recruit anionic phospholipids such as PS to the surface of cancer EVs (**Figure 2G**).

To explore potential reasons for the anionic ζ-potential of Pro-K treated EVs, we further examined the PS level on EVs after Pro-K treatment and observed that the PS was not affected by the Pro-K treatment (**Figure 2H**. The continued anionic ζ-potential of EVs treated with Pro-K was likely due to the continued presence of PS. To evaluate whether anionic phospholipids apart from PS could potentially contribute to the overall negative ζ-potential of EVs, we synthetically fabricated liposomes with increasing percentages of PS and determined the ζ-potential of such liposomes. We found that synthetically fabricated liposomes with 70% PS obtained the largest negative ζ-potential, but still below that of tumorigenic cell derived EVs (**Figure 2I**). Overall, these results indicate that PS is not affected by Pro-K treatment and the negative ζ-potential of Pro-K treated EVs may be due to the continued presence of PS. The results with fabricated liposomes suggest that other anionic lipids or DNA on EV membrane could contribute to the large negative ζ-potential of cancer EVs.

To investigate whether extraluminal DNA on EVs might contribute to the observed negative ζ-potential, the EV surface charge was compared before and after DNase treatment (**Figure 2J-L**). We observed that DNase treatment did not alter the size or the number of the EVs (**Figure 2J**). DNase treatment successfully removed the extraluminal DNA adhering to the EV surface (**Figure 2K**). Interestingly, we observed a drop in the negative ζ-potential of the DNase treated EVs (**Figure 2L**). This result indicates that extraluminal DNA also contributes to the overall negative charge on the EV surface, which is higher in cancer EVs.

### On-chip Microfluidic ElectroPhoresis (MEP) platform enriches anionic EVs

We measured the ζ-potential of EVs extracted from the serum of PDAC patients and healthy donors and found no significant differences between their ζ-potential (**Figure 3A**). Since large negative ζ-potential is a unique property of PDAC cell line derived EVs (**Figure 1A, I**), we hypothesized that cancer EVs may constitute a much smaller fraction of total serum EVs. Serum-derived EVs in cancer patients are a heterogenous mix of cancer and normal tissue-derived EVs, varying by patient. To assess whether cancer and non-cancer EVs in patient serum exhibit distinct ζ-potentials, as seen in cell lines, we needed a method to enrich EVs based on their charge, as standard isolation techniques do not distinguish ζ-potential. To achieve this goal, we developed a microfluidic electrophoresis (MEP) platform to enrich for more anionic EVs (**Figure 3B**).

**Figure 3:**
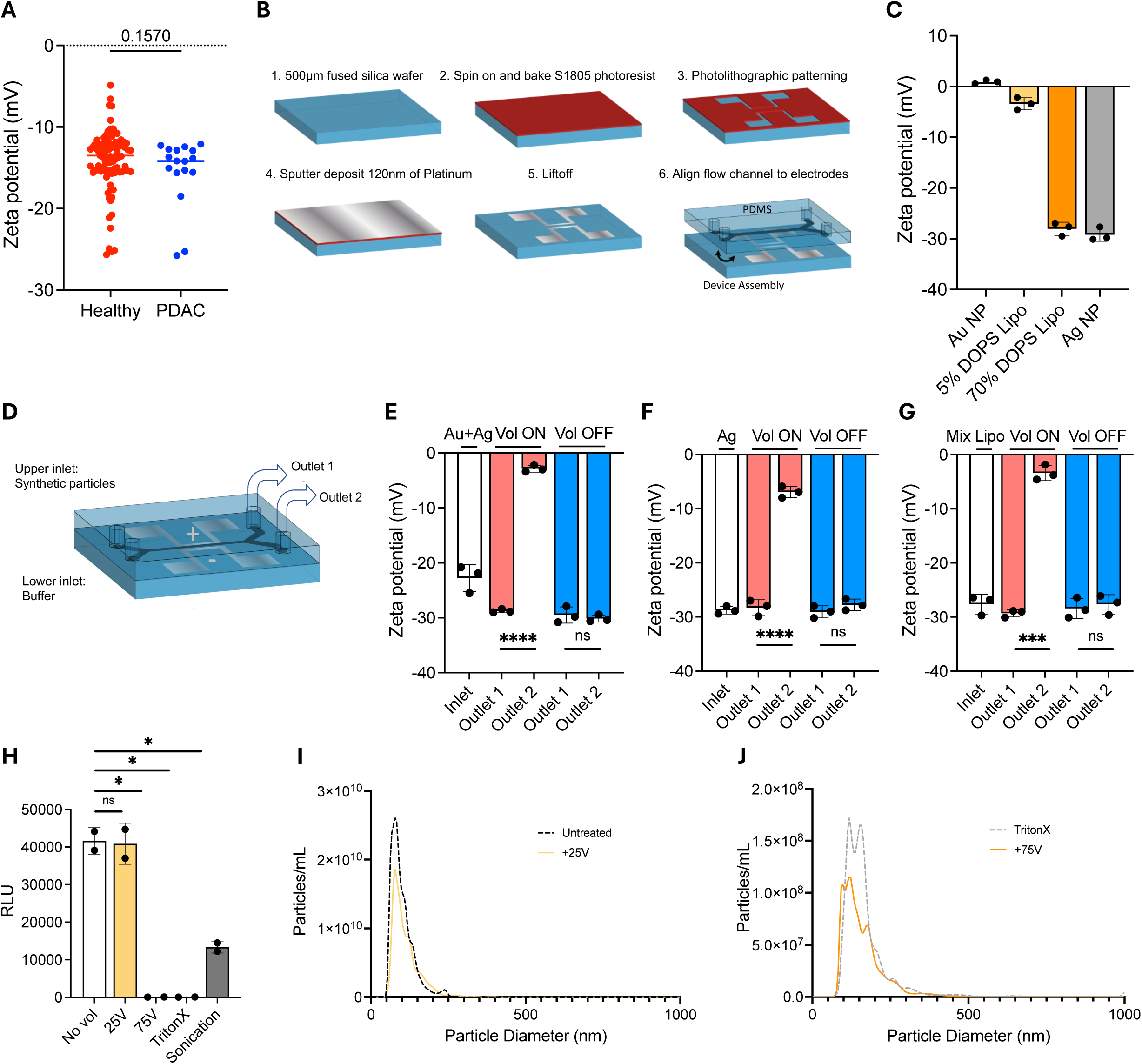
An on-chip Microfluidic ElectroPhoresis (MEP) platform designed to enrich anionic synthetic particles. **(A)** ζ-potential of EVs from PDAC and healthy serum quantified by Malvern Zetasizer (n = 3 technical replicates). **(B)** Schematic of the fabrication process of the MEP chip. Layout for the MEP chip. **(C)** ζ-potential of silver (Ag) nanoparticles, gold (Au) nanoparticles, and anionic liposomes with 5% or 70% DOPS, respectively, quantified by Malvern Zetasizer (n = 3 technical replicates). **(D)** Design of the system for enrichment of high anionic synthetic particles. The charged particles and buffer are loaded via upper inlet and lower inlet, respectively, and an electric field is applied across the horizontal microfluidic channel. **(E-G)** ζ-potential showing selective enrichment of mixed Ag and Au nanoparticles, enrichment of pure highly anionic Ag nanoparticles, enrichment of high anionic liposomes (70% DOPS + 30% DOPC) respectively, quantified by Malvern Zetasizer (n = 3 technical replicates). **(H)** Luminescence intensity of EVs with membrane-anchored luciferase after specific treatment. **(I-J)** Size characterization of EVs after specific treatment. Data are represented as Mean ± SD. Statistical significance was determined using Unpaired t-test (∗p ≤ 0.05, ∗∗p ≤ 0.01, ∗∗∗p ≤ 0.001, ∗∗∗∗p ≤ 0.0001,) n = 3 biological replicates unless specified.

We validated the effectiveness of the newly designed MEP platform to separate based on charge using synthetic nanoparticles. The synthetic nanoparticles employed for validation were cationic gold (Au) nanoparticles (NPs), Silver (Ag) NPs and liposomes (5% DOPS and 70% DOPS). Liposomes with 5% DOPS had a charge near neutral, −4.3±1.4mV, and liposomes with 70% DOPS were highly anionic at −30.1±2.1mV (**Figure 3C**). The synthetic nanoparticles suspension and buffer were loaded via upper and lower inlets respectively and an electric field was applied across the horizontal microfluidic channel (**Figure 3D**). The separation was quantitatively assessed by measuring the ζ-potential of the recovered EVs from both outlets at both voltage-on and voltage-off conditions (**Figure 3E-G**). In the voltage-on condition, more anionic particles were recovered from outlet 1. However, in the voltage-off condition no significant differences in the ζ-potential of the recovered outlets were observed for the different synthetic nanoparticles employed here. The results suggest an enrichment of negative particles in outlet-1 due to the electrophoretic properties.

### Evaluation of EV integrity after exposure to electric field

For electrophoresis-driven enrichment, the application of an electric field may damage EVs and compromise their surface membrane integrity. In this study the electric field EVs experience is approximately 330 V/cm, which yields a transmembrane potential (V_tm_) of 5 mV for an average 100 nm EV. This transmembrane potential is lower than the potential required to induce electroporation (10 mV) or to cause cellular lysis (1 V). While the applied voltage in the MEP platform did not appear to affect the EVs, as an additional validation, we tested the integrity of EVs after application of the electric potential by employing EVs that were tagged with membrane-anchored luciferase with their active domain exposed on the surface. EVs were subjected to a 25 V pulse and a 75 V pulse, and the luminescence intensity was measured (**Figure 3H**). As a control, sonicated or lysed (TritonX treatment) EVs were used to simulate the membrane rupture. In the case of the positive control, overall luminescence decreased after rupture, lysis, or under 75 V pulse but remained unchanged in the 25 V pulse applied or untreated groups (**Figure 3H**). Size distribution determined by NTA before and after application of the 25 V pulse remained unchanged (**Figure 3I**), indicating the integrity of the EV surface membrane along with the unaltered luminescence intensity; however, under the 75 V pulse or TritonX treatment condition, variation in the size distribution and decrease of particle number were observed along with a decrease in luminescence intensity, which is suggestive of EV membrane damage (**Figure 3I-J**). The experiments here also suggest that the voltages that could induce membrane rupture as employed during electroporation are much higher than that used in the MEP device here. Overall, these results indicate that the integrity of EVs are not compromised by the voltage applied for EV enrichment using the MEP platform.

### Enrichment of cancer EVs using MEP platform

After validating the platform with synthetic nanoparticles, we examined the ability of the MEP device to enrich EVs with specific ζ-potential features. Panc1 EVs which exhibited the highest negative ζ-potential (**Figure 1A**), were loaded via the upper inlet at a rate of 5.5 μL/min, and buffer (EV diluent only) was loaded through the lower inlet at a rate of 11 μL/min. An electric field was applied across the horizontal microfluidic channel. To confirm that large negative ζ-potential EVs migrate preferentially towards a positive electrode region, a voltage of 25 V was applied. EV concentration of both inlet and the recovered outlets for the voltage-on/off condition was measured to evaluate for enrichment of cancer EVs (**Figure 4A**). The concentration of EVs recovered from the outlet 2 in the voltage-on condition was in the lower range of 10^7^ EVs/mL whereas particles recovered from outlet 1 was in a range of 10^10^ EVs/mL, a significantly higher concentration than that of outlet 2. The NTA profile of outlet 1 recovered EVs were more representative of the small EVs when compared with outlet 2 recovered EVs in the voltage-on condition. In the voltage-off condition, the inlet EV concentration was in the range of 2 x 10^11^ to 3 x 10^11^ EVs/mL, while the particles recovered from outlets 1 and 2 had concentrations ranging between 8 x 10^10^ to 1 x10^11^ (**Figure 4B**). These results suggests that the MEP platform in the voltage-on condition enriched for anionic cancer EVs via their attraction to positive electrode and recovered from outlet 1.

**Figure 4:**
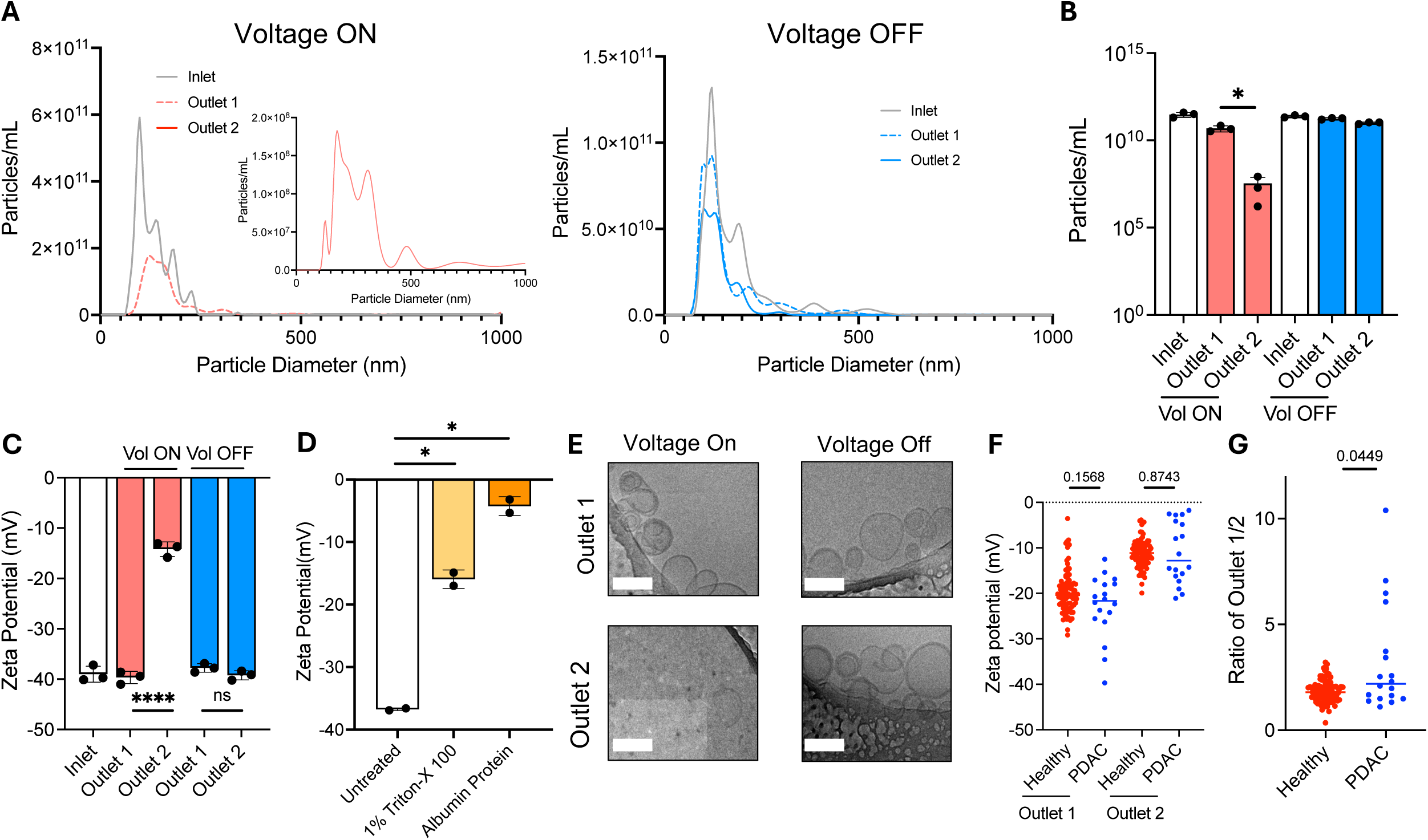
MEP platform enables enrichment of anionic EVs and discriminates PDAC patients from healthy donors. **(A)** Size characterization for inlet, outlet-1 and outlet-2 for voltage-on and off conditions. **(B)** Quantitative characterization of EVs enrichment, showing the concentration of EVs at inlet, outlet-1 and outlet-2 for voltage on and off conditions (n = 3 technical replicates). **(C)** ζ-potential of inlet and particles recovered from outlets 1 and 2 in voltage-on/off condition, quantified by Malvern Zetasizer. **(D)** ζ-potential of untreated EVs, Triton-X treated EVs, and albumin, quantified by Malvern Zetasizer. **(E)** Cryo-EM images of EVs showing integrity from recovered outlets, post passage through the device in voltage-on/off condition. Scale bars, 100 nm. **(F)** ζ-potential of particles recovered from outlets 1 and 2 for voltage on conditions quantified by Malvern Zetasizer (n = 3 technical replicates). **(G)** Ratio of ζ-potential of particles recovered from outlets 1 and 2 in voltage on condition (n = 3 technical replicates). Data are represented as Mean ± SD. Statistical significance was determined using Unpaired t-test (∗p ≤ 0.05, ∗∗p ≤ 0.01, ∗∗∗p ≤ 0.001, ∗∗∗∗p ≤ 0.0001,) n = 3 biological replicates unless specified.

Next, we used zeta sizer to determine the ζ-potential of the recovered EVs from the outlets. In the voltage-off condition, insignificant difference between outlets 1 and 2 was detected (**Figure 4C**). However, in the voltage-on condition, there was a significant increase in ζ-potential of the EVs recovered from outlet 2 compared to EVs recovered from outlet 1, suggesting the enrichment of higher anionic EVs in outlet 1 (**Figure 4C**). The negative ζ-potential in outlet 2 recovered EVs in the voltage-on condition could be attributed association with random proteins and other biomolecules associated with lower ionic EVs. These potential contaminants may precipitate along with EVs in the pellet when isolating EVs by ultracentrifugation isolation technique. EVs usually acquire the negative surface charge from the anionic phospholipid membrane, whereas the proteins and other biomolecules acquire their electric charge from side-chain amino acids or ionizable groups and they usually possess an overall negative ζ-potential that is lower in magnitude than EVs^40^.

Albumin (BSA) is the most abundant protein in the blood and lysed EVs. To exclude the influence on the ζ-potential from BSA, we next evaluated the ζ-potential of ultracentrifuge-isolated EVs, lysed EVs, and BSA. We found that intact cancer EVs possessed significantly higher negative ζ-potential (−36.9 ± 0.3 mV) compared to membrane ruptured cancer EVs (−15 ± 2 mV) or BSA (−5.34 ± 2.13 mV) (**Figure 4D**). These results indicated that the recovery efficiency of EVs with high negative ζ-potential was much higher in outlet 1 in the voltage-on condition.

Cryo-EM was performed to further characterize the particles recovered from outlets 1 and 2 for voltage-on/off conditions (**Figure 4E**). A large number of particles ranging in size from 80-150 nm were detected in outlet 1 in the voltage-on condition with morphology consistent with EVs (**Figure 4E**). Cryo-EM did not detect any EV like structures in the samples collected from outlet 2 in the voltage-on condition; while in the voltage-off condition, EVs were detectable in both outlets by cryo-EM (**Figure 4E**). Collectively, these results confirm that the MEP platform is capable of enriching EVs with negative ζ-potential in outlet 1 due to electrophoretic properties.

Based on the conditions optimized for the MEP platform using synthetic nanoparticles and cell-line derived cancer and normal EVs, we proceeded to test the new MEP device for EVs derived from PDAC patient serum samples to determine whether MEP platform could enrich cancer EVs from serum. EVs were isolated from the serum of healthy and PDAC donors. Serum EVs from the PDAC patients or healthy donors were fractionated using the MEP platform and the size of EVs before and after the enrichment using the MEP platform was evaluated (**Supplementary Figure 2**). ζ-potential was determined for EVs recovered from the two outlets (**Figure 4F-G**). In the voltage-on condition, the ζ-potential of the EVs recovered from the two outlets revealed more anionic EVs elution via outlet 1 compared to outlet 2, while a more negative ζ-potential was detected in the EVs from serum of PDAC patients (**Figure 4F**). Next, enrichment analysis was performed by determining the ratio of ζ-potential for EVs recovered from outlets 1 and 2 in the voltage-on condition. This ratio of EVs derived from serum of PDAC patients and healthy donors showed an enrichment of more anionic particles in the PDAC serum EV population compared to the healthy serum EV populations (**Figure 4G**). This result suggests the ability of MEP platform to enrich the more anionic cancer EVs from the PDAC serum samples that contain both tumor derived and normal tissue derived EVs. Our ultimate goal of enriching cancer EVs from a pool of PDAC serum EVs via variations in the ζ-potential between cancer EVs and non-cancer-EVs was achieved in this proof-of-concept study using the MEP device.

## Discussion

The potential of using extracellular vesicles and exosomes as biomarkers in cancer detection has not been fully realized because current isolation techniques cannot reliably and rapidly distinguish between cancer EVs and non-cancer EVs from circulation. The existing immunoaffinity-based methods for specific EV selection require specific antibodies/labels, and such markers for isolation do capture all EVs and contaminants, thus producing low or undetermined EV-enrichment ratios due to inherent heterogeneity. Our work addresses these key challenges associated with cancer EV enrichment. We developed a label-free EV enrichment platform and demonstrated the feasibility of enriching EVs based on charge. Here, we have shown differential electrical properties of normal and cancer cell derived EVs, showing that PaCa cell derived EVs possess a larger negative net charge compared to non-cancer cells derived EVs.

Our findings about surface charge of EVs derived from both cell lines and serum, support previous findings for different cancer cell derived EVs^41^ and body fluids^42^. Akagi et al. demonstrated that EVs secreted from six cancer cell lines possessed a large negative charge compared to the non-cancerous counterparts^38^. Several studies^43–51^ have shown that EVs carry an overall negative ζ-potential; therefore, we would not expect EVs to interact with negatively-charged cell membranes, yet such interactions have been documented. One possible explanation for such EV-cell interaction is that cell membranes are asymmetric and have a non-uniform lipid distribution. Some regions of the membranes are rich in anionic or cationic lipids. A cationic lipid, sphingosine, has been shown to involved in the internalization of anionic DNA through electrostatic interaction^43, 52–56^. This suggests that the negatively charged EVs are attracted to cationic microdomains on membrane surface and may succeed in crossing the lipid membrane barrier and get internalized in the recipient cells.

Various biomolecules on the EV surface may contribute to the negative surface charge. As an anionic biomolecule, we investigated whether extraluminal DNA affected the negative surface charge on EVs. Reports have identified extraluminal DNA on the EV surface^57, 58^. In this study, we show that the extraluminal DNA located on the EV surface contributed to the overall anionic EV ζ-potential. We also evaluated the role of Phosphatidylserine (PS), a negatively charged phospholipid, which is preferentially localized in the normal cells to the inner leaflets of the cellular membrane, whereas tumor cells have PS exposed to their outer leaflet of cell surfaces. Interestingly, it has been reported that, EVs that get secreted from cancer cells have PS externalized on the outer leaflets and suggested to be associated with cancer progression^50, 59–65^. Here, we observed that the PS levels measured on the surface of EVs secreted by tumorigenic pancreatic cancer cells were significantly higher compared to EVs secreted from non-tumorigenic pancreatic cells. Moreover, higher negatively charged EVs showed increased levels of PS externalization on their membranes. Both our study and another recent study^42^ found that PS remained unaffected by Pro-K treatment, suggesting that anionic ζ-potential Pro-K treated EVs may result from anionic lipids such as PS.

The contribution of mutant KRAS in EV secretion and charge has been a focus of ongoing research^24^. We assessed the contribution of mutant Kras^G12D^ in EV charge and secretion and found that the EVs secreted from Kras-on cells were significantly more anionic than EVs secreted from Kras-off cells. Interestingly, we observed that EVs secreted from Kras^G12D^ knockdown cells had a significantly lower PS level, resulting in reduced anionic ζ-potential. We also found that mutant KRAS induction resulted in an increase in EV secretion from these cell lines. Likewise, a prior study has shown that the induction of mutant KRAS in colon cancer cells significantly increases EV secretion^24^. Proteomic analysis performed on EVs secreted from mutant vs. wild-type KRAS cells identified several critical differences in their protein content. Researchers have found that mutant KRAS cell derived EVs contained a higher amount of tumor-promoting proteins, such as KRAS, SRC family kinases, integrins, and EGFR^24^. Previous studies have shown that KRAS has a single farnesyl chain preceded by a polybasic domain (PBD) of six lysine residues^66^. This cationic polybasic domain allows KRAS to interact with anionic phospholipids in the plasma membrane through electrostatic interactions^56, 66–68^, resulting in an overall negative surface charge for cells. In our study, we observed a similar connection where EVs from Kras-on cells possessed a significantly larger negative ζ-potential and higher PS level on their surface when compared to EVs derived from Kras-off cells. Furthermore, in case of cells it has also been reported that positively charged polybasic domains of KRAS protein near the plasma membrane attract anionic phospholipids other than PS, such as PIP2, and PIP3, resulting in net negative surface charge on the cellular membranes^66–70^.

One of the hallmarks of the most common pancreatic cancer (PaCa), pancreatic ductal adenocarcinoma (PDAC), is the presence of large intra-tumoral heterogeneity^4, 71^, which makes the disease difficult to diagnose early. Current biomarkers such as CA19-9 do not have sufficient sensitivity^4, 72–74^. Hence to realize the usage of EVs in early diagnosis of PDAC, we developed a novel platform that has the ability to enrich tumor EVs (TEV) by exploiting the electrophoretic mobility difference between cancer cell derived EVs and normal cell derived EVs. To prove this concept, we used clinical samples to validate the ability of the MEP platform to enrich TEV from circulating EVs and to reliably detect for variations in the ζ-potential between the TEV and non-TEV from the real-world clinical samples. The search for consistent stand-alone biomolecular markers found on the EV surface has proven to be very difficult^12, 75, 76^. Further investigation is needed to determine whether utilizing a combination of the biophysical properties of TEV along with the biomolecular properties proves to be a more reliable approach for early detection. In summary, our work provides a novel approach for the enrichment of cancer cell derived EVs based on anionic charge, with potential application for early detection of pancreatic cancer.

## Methods and Materials

### Cell lines

### Cell culture

Tumorigenic and non-malignant cells used for the study are listed in the **Table 1**. The human cell lines used in this study were: Panc-1; MiaPaCa-2; T3M4; BxPC3; HPDE; HPNE; and HK-2. The metridia membrane luciferase expressing MDA-MB-231; CD63 GFP labeled Panc1 cell line derived EVs were used for integrity assay. Panc1, T3M4, BxPC3, HPNE and CD63 GFP+ Panc1 cells were cultured in RPMI 1640 (Corning) supplemented with 10% FBS and 1% penicillin-streptomycin. The primary murine cells (iKras) were isolated from p48 cre; R26-rtTa-IRES-EGFP; TetO-KrasG12D p53L/L (PiKP) PaCa genetically engineered mouse model (GEMM)^77^ and maintained in RPMI-1640 (Corning), 10% Tet System Approved FBS (Clontech 631106), 1X Penicillin-Streptomycin (Corning 30-002-Cl) in the presence of 1 μg/ml doxycycline to ensure oncogenic Kras expression (Kras^G12D^ on cells). To extinguish oncogenic Kras, doxycycline was withdrawn from the media (Kras^G12D^ off cells). HPDE cells were cultured in 50% DMEM (Corning) supplemented with 10% fetal bovine serum (FBS), 1% penicillin-streptomycin and 50% defined keratinocyte-SFM (Gibco) with 1% penicillin-streptomycin. MDA-MB 231 pMetLuc membrane reporter cells (metridia membrane bound luciferase) and HK-2 cells were cultured in DMEM (Corning) supplemented with 10% fetal bovine serum (FBS) and 1% penicillin-streptomycin. Coelenterazine (Fisher Scientific) was used as the substrate of the MDA MB-231 pMetLuc cells and luminescence intensity was measured after 2 min at room temperature using a luminometer (BMG Labtech) Omega Microplate reader. All cell lines were tested for mycoplasma contamination and were maintained in humidified cell culture incubators at 37°C and 5% CO_2_.

**Table 1:** Cell lines.

| Cell line | Growth conditions | Source | Malignant |
| --- | --- | --- | --- |
| Panc1 | RPMI- 10% FBS,1% PS | ATCC | Yes |
| MiaPaCa2 | DMEM- 10%FBS,2.5% Horse<br>Serum,1% PS | ATCC | Yes |
| T3M4 | RPMI- 10% FBS,1% PS | Riken Bioresource | Yes |
| BxPC3 | RPMI- 10% FBS,1% PS | ATCC | Yes |
| HPDE | 1:1 Keratinocyte serum-free<br>media:DMEM, 10% FBS, 1%PS | MDACC, | No |
| HPNE | RPMI- 10% FBS,1% PS | ATCC | No |
| HK-2 | DMEM 10% FBS, 1% PS | ATCC | No |
| iKras-on (Murine<br>primary Kras<sup>G12D</sup> On) | RPMI 10% Tet-FBS, 1% PS | MDACC | Yes |
| iKras-off (Murine<br>primary Kras<sup>G12D</sup> Off) | RPMI 10% Tet-FBS, 1% PS | MDACC | Yes |
| MDA MB-231 pMetLuc<br>membrane reporter | DMEM 10% FBS, 1%PS | MDACC, | Yes |
| Panc1 CD63 GFP+ | RPMI- 10% FBS,1% PS | MDACC, | Yes |

### Generation of pMetLuc and CD63 GFP+ cells

MDA-MB-231 cells were transfected with pMetLuc-Mem Control (Clontech) with Lipofectamine 2000 according to manufacturer’s instructions. Cells were selected with 2 mg/mL G418 (Gold Biotechnology) to establish a stably transfected line. Panc1 (1×10^3^ cells) were transduced in a 96 well plate with 1 µL of pCT-CD63-GFP lentivirus (SBI CTYO12-VA-1). Cells were then selected with 5 µg/mL puromycin (Sigma-Aldrich) to establish a stable line.

### Cell line derived EV production

Cells were grown in T225 flasks to 85% confluence and were washed two times with 1X phosphate-buffered saline (PBS). Then the cells were incubated for 48 h in serum free media. The conditioned media (CM) were centrifuged for 5 min at 800 x g and then for 10 min at 2,000 x g to remove cells and smaller debris. The supernatant was filtered through a 0.2 μm pore filter (Corning, Cat. No. 431219) and then subjected to ultracentrifugation (UC) (Beckman) at 100,000 x g for 3h at 4^◦^C in a Beckman, SW 32 Ti rotor. The supernatants were discarded and the pellet containing EVs were resuspended in 1X PBS for downstream processing.

### Patient samples and tissue collections

Serum samples from healthy participants were obtained as deidentified, discarded clinical material and were therefore exempt from Institutional Review Board (IRB; PA14/0A32) approval. Informed consent was obtained from all participants in accordance with institutional guidelines. Blood samples of healthy individuals were collected by MD Anderson Blood Bank. Samples were extracted after centrifuge at 3,000 x g for 20 min and stored at −80^◦^C until analyzed.

Serum samples from patients diagnosed with PDAC were collected at University of Cologne and University Hospital Heidelberg by the local IRB (Cologne: 13-091; Heidelberg: 323/2004) and written informed consent was obtained from all participants prior to surgery in accordance with institutional guidelines. The information of treatments and stage of tumor is listed in **Supplementary Table 1**. Blood samples were obtained preoperatively, prior to any surgical intervention. Samples were extracted after centrifugation at 2,500 x g or at 4,000 rpm for 10 min and stored at −80^◦^C until analyzed.

A total of 96 healthy participants and 19 PDAC patients were included in this study, with a gender-balanced distribution.

### Human serum-derived EV production

500 μL of serum sample per PDAC patient and healthy donor (n = 112 total) were rapidly thawed at 37°C in a water bath. The serum was centrifuged at 4°C at 500 x g for 5 min, and 2,000 x g for 10 min, and filtered in a syringe with a 0.22 μm membrane filter directly into small ultracentrifuge tubes (Beckman Coulter). PBS was added to the ultracentrifuge tube to fill up to 11 mL. Samples were ultracentrifuged in a SW 40 Ti swinging bucket rotor at 100,000 x g for 3h at 4°C. In the wash step for serum EVs, the EV pellet was resuspended in PBS, the ultracentrifuge tubes were then filled to the top with PBS and samples were ultracentrifuged again in a SW 40 Ti swinging bucket rotor at 100,000 x g for 3h at 4°C. Supernatant was aspirated and pellets were resuspended in 500 μL PBS and stored at −80°C until further use.

### Nanoparticle tracking analysis

EV size and concentration were analyzed using the LM10 nanoparticle characterization system (NanoSight LM10 Malvern) which was equipped with a 488 nm Blue laser and sCMOS camera (NTA, Malvern Panalytical). Aliquots of EV suspension were diluted in cell culture grade water for nanoparticle tracking analysis, and tracking was performed at 25°C with a camera level set at 14-15. A similar threshold was used across all samples for consistent recording. The speed of the syringe pump was set at 20. Three 30 sec videos were captured per sample and the average value was used to determine the size range and concentration of nanoparticles.

### Zeta potential (**ζ**-potential) measurement

ζ-potential of EVs was determined with Malvern Zen 3600 Zetasizer. Anionic and cationic liposomes^11^ were fabricated as controls, and 5×10 particles/mL were tested to generate a standard curve followed by measurements for ultracentrifuge derived EVs. A folded capillary zeta cell (DTS 1070) Malvern Paranalytical was used as the zeta cell cuvette to determine the zeta ζ-potential. Three biological and three technical replicates were performed, with settings (Dielectric constant = 80; viscosity = 0.9; RI (Refractive Index) = 1.340; temperature = 25°C) and pH between 5.5 to 5.9.

### Cryo-EM analysis

In order to quantitatively verify the morphology of the EVs and evaluate the integrity post passage through the chip, we employed cryo electron microscopy (Cryo-EM). For cryo-EM, EVs were resuspended in 50 μL of PBS. The vitrification steps were performed at SEA (Shared Equipment Authority) Rice University. Quanti-foil mesh grids were used for the study. Before the vitrification step the grids were air-glow discharged for 120 sec. For the vitrification process, about 3-5 μL of EV samples were applied on to the grid. After blotting the grids were snap frozen into liquid ethane using the FEI Vitrobot system. The frozen grids were then transferred to liquid nitrogen until imaging. Imaging was performed using a JEOL 2010 Transmission Electron Microscope (TEM) with cryo.

### Bead-based flow cytometry analysis of EVs

To evaluate the surface protein marker expression, 5×10^9^ EVs were resuspended in 200 μL PBS in Eppendorf tubes. 10 μL of Aldehyde/sulphate laEVs beads (4% w/v) (Invitrogen) were added to the solution and mixed at room temperature for 15 min. In order to facilitate the bead binding to EVs, 600 μL PBS was added into each vial, vorEVsed and allowed to rotate at 4^◦^C overnight. The following day, 400 μL Glycine (1M) was added to each vial and incubated at room temperature for 1h with rotation. Next, the bead-bound EVs were precipitated at 12,000 rpm at room temperature for 2 min. Next as a blocking step, the EVs binding the beads were incubated for 60 min in 100 μL of 10% BSA. The suspension was then precipitated at 12,000 rpm for 2 min at room temperature (RT) to wash away excess BSA. The EVs attaching to the beads were then resuspended in 40 μL of 2% BSA in 1X PBS, and were split equally into six tubes to stain for EV surface markers (CD81, CD63, CD9) and for control (secondary antibody only, IgG alone and EV alone). The bead-bound EVs were incubated with unconjugated primary antibodies: 1 μL of Mouse anti-human CD63 (BD 556019, diluted 1:2.5 in 2% BSA); Mouse anti-human CD81 (BD 555675, diluted 1:2.5 in 2% BSA); Mouse anti-human CD9 (Sigma SAB4700092, diluted 1:5 in 2% BSA); Isotype control: Mouse IgG1 κ isotype control (BD 555746, diluted 1:2.5 in 2% BSA). For mouse cell line derived EVs, the following antibodies used were: Rat Anti-mouse CD63 (BD 564221, diluted 1:2.5 in 2% BSA); Biotin hamster anti-mouse CD81 (BD 559518, diluted 1:2.5 in 2% BSA); Biotin rat anti-mouse CD9 (BD 558749, diluted 1:5 in 2% BSA); Isotype control: Purified Rat IgG2a, κ isotype control (BD 553927, diluted 1:2.5 in 2% BSA) in 20 μL volume with 2% BSA, and rotated for 1h at room temperature. After incubating with primary antibody for 1h, the bead-bound EVs were precipitated at 12,000 rpm at room temperature for 2 min and the supernatant was discarded. Post incubation with the unconjugated primary antibodies, the bead-bound EVs were washed 3X in 200 μl of 2% BSA, precipitated at 12,000 rpm for 2 min, and the supernatant discarded. Next, the bead-bound EVs were incubated for 1h rotating in 20 μL of 2% BSA containing the secondary antibody (donkey anti-mouse, Alexa Fluor 488 (Invitrogen A21202), 1:20 in 2% BSA). After staining with the secondary antibody, the samples were then precipitated at 12,000 rpm at room temperature for 2 min. Finally, the bead-bound EVs were washed 3X with 2% BSA in PBS. The EV surface markers CD9, CD63, and CD81 on bead-bound EVs were analyzed by LSR Fortessa X-20 (BD Bioscience). Quantification of positive EVs for CD81, CD63, and CD9 was performed in FlowJo Version 10.6.1. EV samples that were collected from outlet post passage through the chip, were washed and ultracentrifuged for 3h at 4^◦^C with a speed of 28,000 rpm in a Beckman, SW 32 Ti rotor. The particles from both outlets were conjugated to the aldehyde sulfate laEVs beads and bead-based flow cytometry was used to quantify EV markers.

### Anchorage-independent growth assay

Soft agar was used to examine the anchorage-independent growth. CytoSelect 96-Well Cell Transformation Assay (Soft Agar Colony Formation Catalog# CBA-130-T) from Cell Biolabs was utilized as per the manufacturer’s protocol. Briefly, 50 μL of the base agar matrix was dispensed into the wells of a 96-well plate. At the bottom surface of each well once the agar was solidified, cell suspension^3^ of 75 μL containing 10×10 cells was layered on top of the agar. 100 μL 2x complete RPMI medium with 10% FBS and 1% penicillin-streptomycin was added on the cell suspension. On the eighth day the colony formation was examined under a light microscope. Next the quantitation of Anchorage-Independent growth was performed by fluorometric detection. Eight days post incubation, in order to solubilize the agar matrix completely, 50 μL of the Agar Solubilization Solution was added to each well and incubated for 1h at 37^◦^C. In order to ensure the complete agar solubilization each well was pipetted 5-10 times. Next 25 μL of the 8X lysis buffer was added into each well and pipetted well to ensure a homogeneous mixture. After incubation of the plate for 15 mins at room temperature, 10 μL of the lysed mixture was transferred to a 96-well plate that was suitable for fluorescence measurement. 90 μL of the CyQuant^©^ GR dye solution was added into each plate and incubated 10 mins at room temperature. When CyQuant^©^ GR dye binds to the nucleic acids it exhibits strong fluorescence enhancement. In order to determine the fluorescence from the colonies, the plate was read in a 96-well fluorometer. The fluorescence produced by each well was proportional to the number of viable cells.

### Cleavage of surface proteins with Proteinase K

EVs isolated by ultracentrifugation were used. For the preparation of Proteinase K (Pro-K) - digested EVs - purified EVs were incubated with 5 mg/mL of Proteinase K (dissolved in RNase-free water Sigma-Aldrich P2308) at 37^◦^C for 30 min. This was followed by an inactivation step were ProK-treated EVs were heated at 50^◦^C for 20 min. EVs were washed post Pro-K treatment with PBS at 28,000 rpm for 3h using ultracentrifugation in a SW40 Ti swinging bucket rotor. The washed EVs were collected and concentration was assessed by Nanosight^TM^ (Malvern). The collected EV samples were aliquoted to limit the freeze thaw cycles. The samples were stored at −80^◦^C until use to probe for biomolecular and physicochemical properties. The cleavage of ectodomain of EV surface proteins with Pro-K treatment was validated by flow cytometry. The aldehyde-sulphate bead flow cytometry approach described above was used. EV integrity after ProK treatment was determined by quantifying the release of select luminal proteins. Panc1 cells expressing CD63-GFP in the intra-luminal region were used to extract EVs, and integrity was determined by comparing the GFP levels of EVs before and after treatment by single particle detector on the LSR Fortessa X-20 flow cytometer.

### DNase treatment of EVs

EVs treated with 2 U/μL Turbo DNase (AM2238) in 1X TURBO DNase buffer and incubated for 30 min at 37^◦^C. Turbo DNase was inactivated by incubation in 2mM EDTA and heated at 65^◦^C for 10 min. After DNase digestion, samples were washed in PBS and precipitated by ultracentrifugation at 28,000 rpm for 3h in a SW40 Ti swinging bucket rotor. All washed EVs were collected and concentration was assessed by Nano sight (Malvern). Removal of extraluminal DNA on EVs surface was evaluated by measuring the DNA concentrations before and after the TURBO DNase treatment by Nanodrop spectrophotometer.

### Lysis of EVs

Isolated EVs were lysed with 1% Triton X-100 for 30 min at 37^◦^C with 5X protease inhibitor. The samples after treatment with 1% Triton X-100 were washed with PBS and spun down by ultracentrifugation in a SW40 Ti swinging bucket rotor at 28,000 rpm for 3h at 4^◦^C. Protein quantification of lysed and un-lysed EVs was determined by microBCA assay (Thermo Fisher Cat no. 23235).

### Assessment of phosphatidylserine (PS) levels on EVs by flow cytometry

Similar to the EV bead-flow cytometry protocol mentioned above, briefly 2×10^10^ EVs were resuspended in 200 μL of PBS and 10 μL of Aldehyde/Sulphate laEVs beads (Invitrogen A37304) were added to the EVs, followed by rotation for 15 min at room temperature. 600 μL of PBS was added to the solution and was rotated overnight at 4^◦^C. 400 μL of Glycine was added and rotated for 1h at room temperature. This mixture was precipitated at 12000 rpm at RT for 2 min. The pellet was resuspended in 100 μL of 10% BSA and mixed for 1h. The mixture was centrifuged at 12,000 rpm for 2 min at RT and the supernatant was discarded. The bead-bound EVs were resuspended in 40 μL of 2% BSA in PBS, and split equally into two tubes: for staining APC annexin V and for control (EV alone unstained). EVs were stained with APC Annexin V (Cat no. 640941 BioLegend, 1:20) diluted in 1x Annexin V binding buffer rich in calcium ions at room temperature for 1h in the dark. The bead-bound EVs were washed 3 times with 1x Annexin binding buffer in PBS and analyzed for Annexin V using the LSR Fortessa X-20-BD Bioscience. As a control, beads were also stained with APC Annexin V BioLegend (Cat no. 640941) diluted in 1x Annexin V. Quantification of positive EV for Annexin V was performed in FlowJo Version 10.6.1 and overtone levels were determined.

### RT-qPCR for the iKras cell lines

For cell culture, Trizol Reagent (Invitrogen #15596026) was directly added to cell culture plate and total RNA was extracted using Direct-zol RNA MiniPrep Kit (Zymo Research #R2052). 2 μg of RNA was used for cDNA synthesis by High-Capacity cDNA Reverse Transcription Kit with RNase Inhibitor (Applied Biosystems #4374966). qPCR was run with Fast SYBR Green Master Mix (Applied Biosystem #4385612) and the primer sets described in **Table 2**, on the QuantStudio 7 Flex Real-Time PCR System. Transcripts were normalized to 18S levels. The relative fold change expression was used to express the results.

**Table 2:**
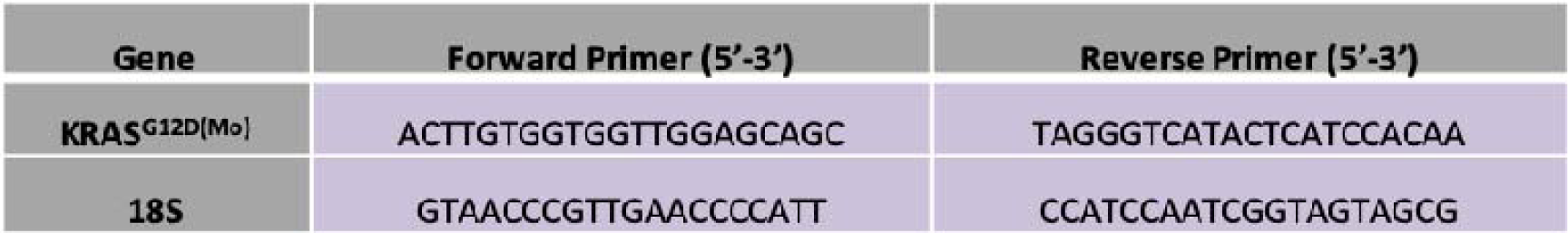
Primer sets for RT-qPCR.

### MEP device fabrication

Flow channel and electrode channel photomasks were designed in K-Layout Version 0.25.9 and printed by Out-put city CAD services. For fabricating electrode wafer, a 500μm fused Silica wafer was plasma cleaned for 30 sec. S-1805 was spin coated and baked at 110°C for 1min. For photolithographic patterning exposure of 95 mJ/cm^2^ was used to pattern the electrode, followed by soaking in MF 321 for 120 sec with gentle agitation. The wafer was rinsed in DI water and dehydrated for 5min at 95°C followed by plasma cleaning for 30 sec. A 120 nm thin layer of platinum was sputter deposited with parameters: Pt, 25W, 3mT, Ar, 35sccm for 60 min. After sputter coating an acetone liftoff for 1h was performed on the shaker. The wafer was rinsed with DI water and dried under N_2_ air. The electrode wafer was then stored in a dry room temperature environment. For fabricating the flow channel, SU-8 3025 was spin coated on the wafer to achieve a target height of 20 μm followed by a prebake for 14 min at 95°C. The flow channel pattern was exposed at 355 mJ/cm^2^ and post-baked at 65°C for 5min followed by ramping at 10 C/min for 10 min at 95°C. The wafer was developed in a SU8 developer for 8.5 min, rinsed with IPA and dried using N_2_ gas. Finally, the flow channel wafer was hard-baked at 65°C for 5min followed by ramping at 10 C/min for 20 min at 150°C. Once polydimethylsiloxane (PDMS) (Sylgard, Dow Corning; 10:1 elastomer: cross-linker weight ratio) was cured on the flow channels, it was peeled off. Holes were punched to connect the tubing. The flow channels and the electrode wafer were aligned, pressed down, and thermally bonded at 80°C for 16h. The completed device was then soldered to wires and the solder connections were insulated using JB-weld epoxy.

### Device preparation

Prior to flow, devices were functionalized overnight with 10% BSA to avoid non-specific adsorption of EVs due to the hydrophobic nature of PDMS (See supplementary Figure 4C, D). Ultrasensitive quartz crystal microbalance (QCM-D, Q-Sense E4 System, Biolin Scientific) was also utilized to confirm no interaction of EVs with BSA layer. The QCM-D sensor surface was coated with BSA and EVs were channeled through the surface but no significant change in frequency (mass adsorption) was observed (see supplementary Figure 4E). For the voltage-on experiments voltage was applied across the electrodes at 25V by a DC power supply (Mastech HY3005D). The inlet 1 was the exosomal sample inlet and EVs solution was passaged at a rate of 5.5 μL/min and inlet 2 was the buffer inlet and buffer (PBS) was passaged at a rate of 11 μL/min (New Era Pump Systems NE-300).

### Synthetic nanoparticles and liposomes experiments

1,2-dioleoyl-sn-glycero-3-phospho-L-serine (18:1 PS (DOPS)), 1,2-dioleoyl-sn-glycero-3-phosphocholine (18:1 (Δ9-Cis) PC (DOPC)) and 1,2-dioleoyl-3-trimethylammonium-propane (18:1 TAP (DOTAP)) from Avanti were used. Different ratios of lipids were mixed in 1 mL chloroform and then dried with N_2_ gas. The resulting lipid films were place in a vacuum chamber for 1h to evaporate the chloroform. Then, the lipid films were hydrated by 1 mL PBS followed by vigorous mixing. The liposomes were extruded with 100 nm diameter polycarbonate membrane for 15 times. The bulk liposomes solution was stored at 4°C.

### Western blot

After running the EV samples through the chip, the collected outlets were washed and ultracentrifuged for 3h at 40,000 rpm in SW40 Ti swinging bucket rotor. To determine the protein levels, the particles recovered from the upper and lower outlets were lysed using a solution of Urea buffer (8M urea, 2.5% SDS, 1 mM phenylmethylsulphonyl fluoride and 1 ug/mL complete protease inhibitor cocktail (Sigma Aldrich Roche 11697498001)). The lysates were loaded onto pre-cast polyacrylamide gels and transferred to polyvinylidene fluoride (PVDF) membranes using the BioRad Turbo Transfer unit. For the blocking step, membranes were incubated at room temperature for 1h with TBS-T containing 5% BSA, and incubated at 4°C with the primary antibodies for overnight: CD63 (E-12) monoclonal (sc-365604) antibody 1:1000 Santa Cruz Biotech, INC; CD81 (B-11) sc-166029 1:1000 Santa Cruz Biotech, INC. Next day, the excess primary antibody was washed for 10 min 3X with TBS-T. Membranes were incubated in secondary antibodies HRP conjugated (HAF007 R&D Systems 1:1000) for 1h at room temperature. After the secondary antibody incubation, membranes were washed for 10 min 3X with TBS-T. Finally, the blots were developed using the West-Q Pico ECL Solution kit (GenDEPOT, Cat No. W3652-020).

### Statistical analysis

All data are expressed as mean values or standard deviation (S.D.), as indicated. Statistical analysis was performed using GraphPad Prism version 10. Statistical analysis used are specified in detail in the respective figure legends.

## Acknowledgements

This work was supported by funds from Sid W. Richardson Foundation to RK. EV work in the Kalluri lab is supported by NCI R35CA263815. We are grateful to the Shared Equipment Authority staff at Rice University for their support. We are grateful to Dr. Wenhua Guo for training KSK on cryogenic electron microscopy. We appreciate Dr. Valerie S Kalluri and Michelle Kirtley for facilitating the connection and assistance with serum samples.

## Author Contributions

Raghu Kalluri – Idea Generation and Conceptualization, Funding acquisition, Project oversight, Project administration, Resources, Supervision, Writing – original draft

Kshipra S Kapoor – Co-conceptualization, Data curation, Formal analysis, Methodology, Software Validation, data Visualization, Writing – original draft, Writing – review & editing

Xin Luo: Data curation and assessment, Formal analysis, Methodology, Writing – original draft, Writing – review & editing

Kaira A Church: Data curation and assessment, formal analysis, Methodology, Writing – original draft, Writing – review & editing

Martin M Bell – Methodology, Software, Fabrication and Validation experiments Bo Fan – Methodology, Software, Fabrication and Validation experiments Seoyun Kong – Investigation – assisted KSK.

Elena V Ramirez – Investigation – assisted KSK with FACS experiments and analysis

Yi-Lin Chen – Investigation – assisted KSK with ζ-potential analysis and experiments, fabricated liposomes and performed QCM-D experiments

Fernanda G Kugeratski – Investigation - assisted KSK with FACS experiments and analysis Sibani Lisa Biswal – Resources (Zeta analysis, QCM-D, Liposome fabrication) and supervision

Kathleen M McAndrews - Investigation – assisted KSK with Luciferase assay experiment and provided the blinded serum sample and unblinded the samples.

Florian Gebauer – Resources (Patient samples) Christoph Kahlert – Resources (Patient samples)

Jacob T Robinson – Resources (Device Fab, and related resources) and supervision

---

**Figure Supplementary 1:**
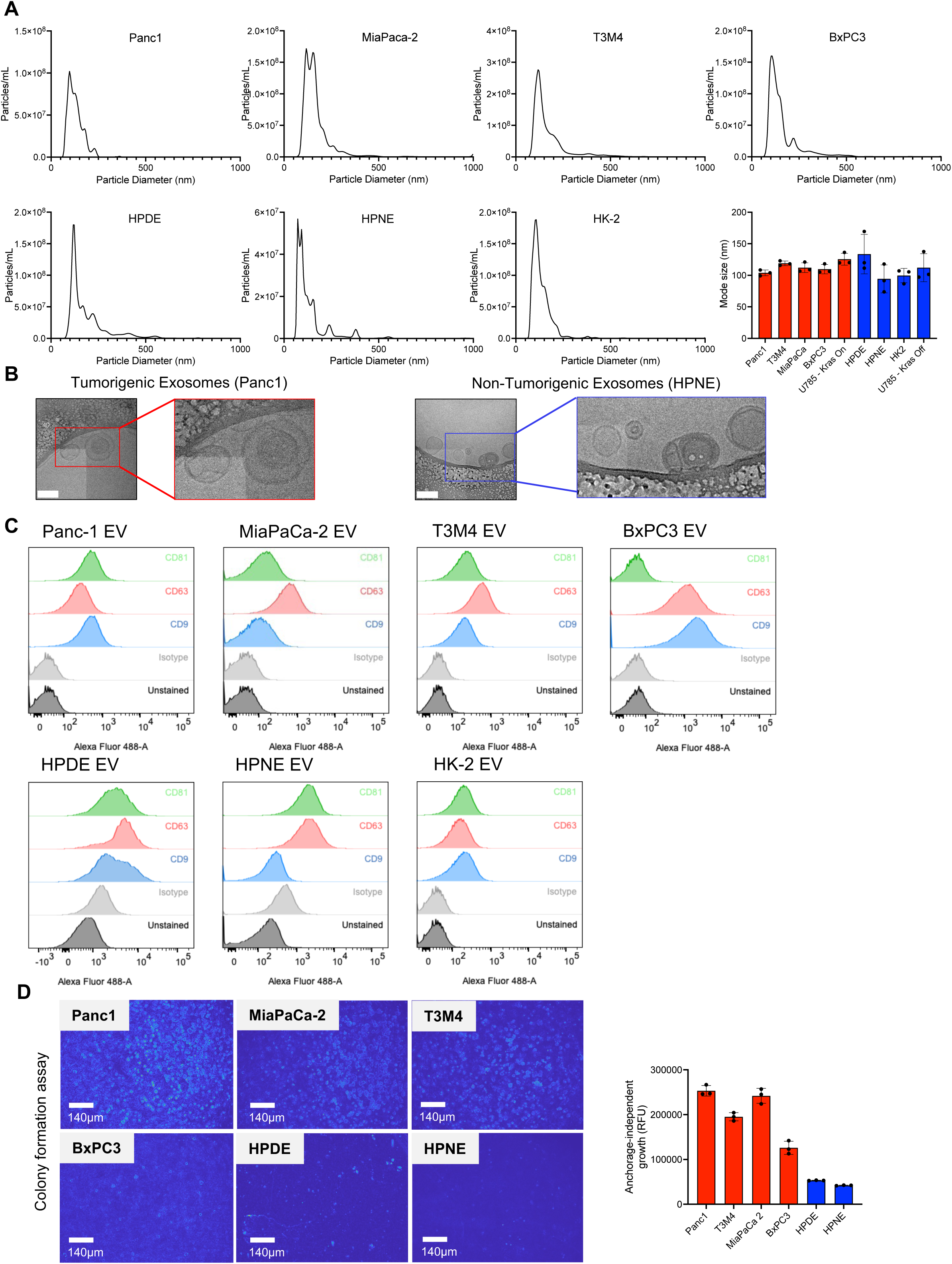
Size and morphological characterization of EVs. **(A)** Particle concentration and size of EVs derived from tumorigenic and non-tumorigenic cell lines. **(B)** Morphology of tumorigenic and non-tumorigenic EVs, determined by Cryo-EM imaging. Scale bars, 100 nm. **(C)** Flow cytometry analysis of EVs for common tetraspanin EV markers with representative histograms for indicated cell lines. **(D)** Aggressiveness of tumorigenic and non-tumorigenic cells tested in soft agar assay. Scale bars, 140 μm. Data are represented as Mean ± SD.

**Figure Supplementary 2:**
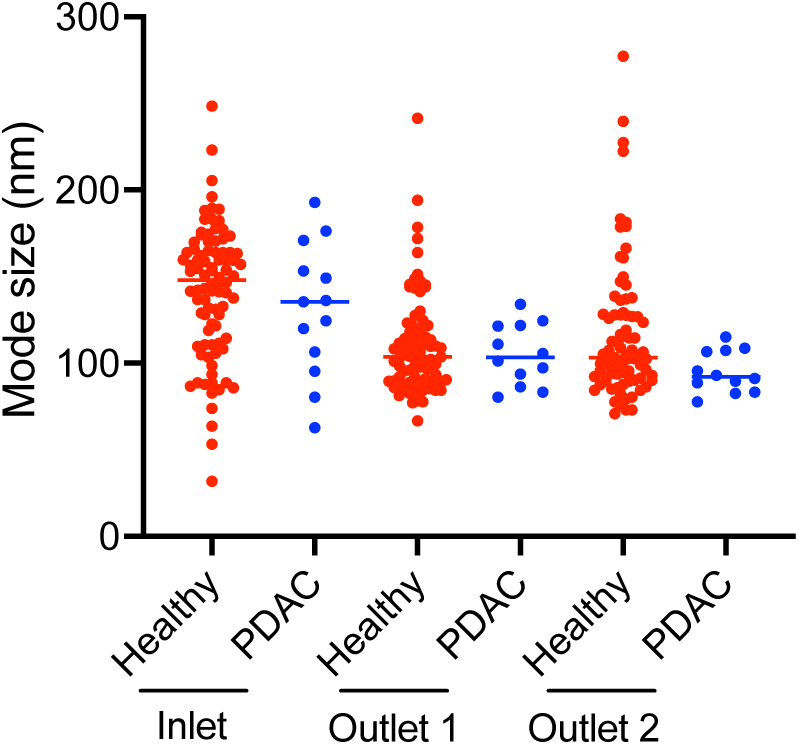
Mode size of EVs from PDAC and healthy donors recovered from inlet and outlets under voltage on condition.

